# SkeleSketch: An Open-Source Tool for Quantifying Microglial Morphology in Fiji/ImageJ

**DOI:** 10.64898/2026.09.24.754118

**Authors:** L.A. Pascoe, V.M. Masegosa, S. Liu, Q. Zhu

## Abstract

Microglial morphology is widely used as an indicator of cellular state, yet existing analytical approaches are often labor-intensive, require proprietary software, or depend on high-resolution imaging platforms. Here, SkeleSketch is presented as an open-source Fiji/ImageJ plugin that automates microglial skeletonization, morphological quantification, and data organization while allowing optional user-guided correction. Using spinal cord microglia from PBS- and LPS-treated mice, SkeleSketch accurately reconstructed cellular morphology and captured expected LPS-induced morphological remodeling, including enlarged soma size and reduced branching complexity. Comparable measurements were obtained from both CX3CR1-GFP reporter mice and Iba1 immunostaining, demonstrating robustness across labeling strategies. Additionally, SkeleSketch morphometrics were positively correlated with those generated by the commercial software Imaris. Our pipeline provides an accessible, reproducible, and flexible platform for quantitative microglial morphology analysis using standard epifluorescence microscopy.

**Significance Statement:** Quantitative analysis of microglial morphology is increasingly used to study how these cells respond to injury, disease, and aging. However, in practice, it remains difficult for many laboratories. Manual tracing is labor-intensive and subjective, whereas automated pipelines typically require high-resolution imaging, expensive software, or computational expertise. SkeleSketch was developed to fill the gap between these approaches. It is a free, open-source Fiji/ImageJ tool that operates on standard epifluorescence images and requires only modest computing resources and minimal technical expertise, with optional user correction where automated tracing struggles. By lowering the technical and financial barriers to reproducible morphometric analysis, SkeleSketch broadens access to quantitative, single-cell microglial phenotyping across basic, translational, and educational neuroscience.

## Introduction

Microglia are the resident macrophages and principal innate immune cells of the central nervous system (CNS), comprising approximately 5-10% of all CNS cells. Beyond their classical roles in immune surveillance, phagocytosis, and maintenance of CNS homeostasis, microglia actively regulate synaptic pruning, adult neurogenesis, and neuronal circuit modulation across the lifespan (Sierra et al., 2010; Saijo et al., 2011; Schafer et al., 2012; Colonna and Butovsky, 2017; Prinz et al., 2021). Alterations in microglial morphology and function have been implicated across a wide range of neurological conditions, including neurodegenerative disease, traumatic injury, infection, and aging, making accurate characterization of microglia important in both basic and translational neuroscience (Salter and Stevens, 2017; Hammond et al., 2019).

Microglial morphology has long served as an accessible indicator of cellular state and function. Traditionally, microglia have been categorized into discrete morphological states, including “ramified” (i.e., resting/surveillant), “amoeboid” (i.e., phagocytic/activated), or rod (Kettenmann et al., 2011). However, these categories fail to capture the broad spectrum of states microglia may occupy (Paolicelli et al., 2022; Green and Rowe, 2024). Disease stage, anatomical region, chronic stress, aging, and biological sex can influence morphology and produce intermediate states with overlapping structural and functional characteristics (Calcia et al., 2016; Masuda et al., 2020). Consequently, assessment of a range of morphometrics (e.g., soma size, branch length, branching complexity) is increasingly recognized as a more objective and informative approach than categorical classification alone.

Despite its widespread use, quantitative microglial morphology analysis remains technically challenging. Manual tracing provides high user control and can accurately capture complex architecture, but is labor-intensive, time-consuming, and subject to user variability, as the tracer reconstructs the entire skeleton de novo (Young and Morrison, 2018). More recent computational approaches have substantially improved throughput and analytical sophistication, but require high-resolution images (e.g., confocal, two-photon), specialized computational expertise, or proprietary platforms (Table 1), which are not accessible to all research groups. Even recent open-source, Fiji-based tools such as MicrogliaMorphology (Kim et al., 2024) can require more technical setup to use and depend on global automated thresholding, making them particularly sensitive to image quality. Consequently, a gap remains between simple but labor-intensive manual approaches and highly sophisticated computational pipelines that are not easily adopted in routine laboratory workflows.

**Table 1.**
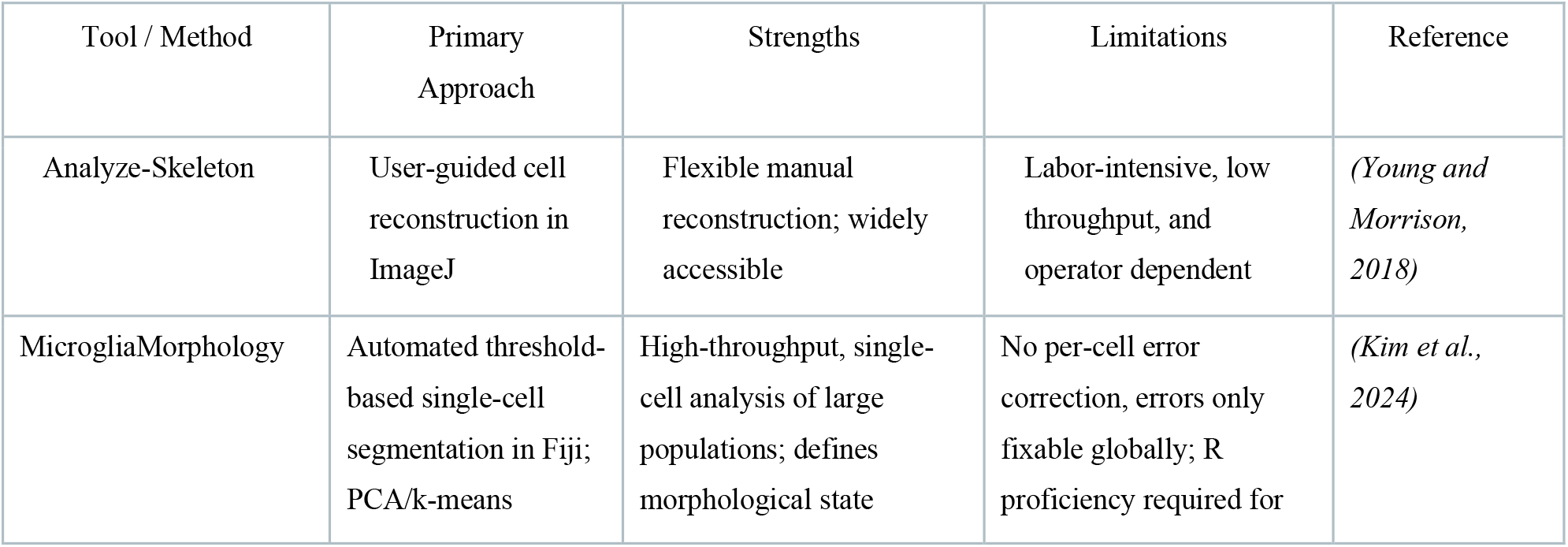
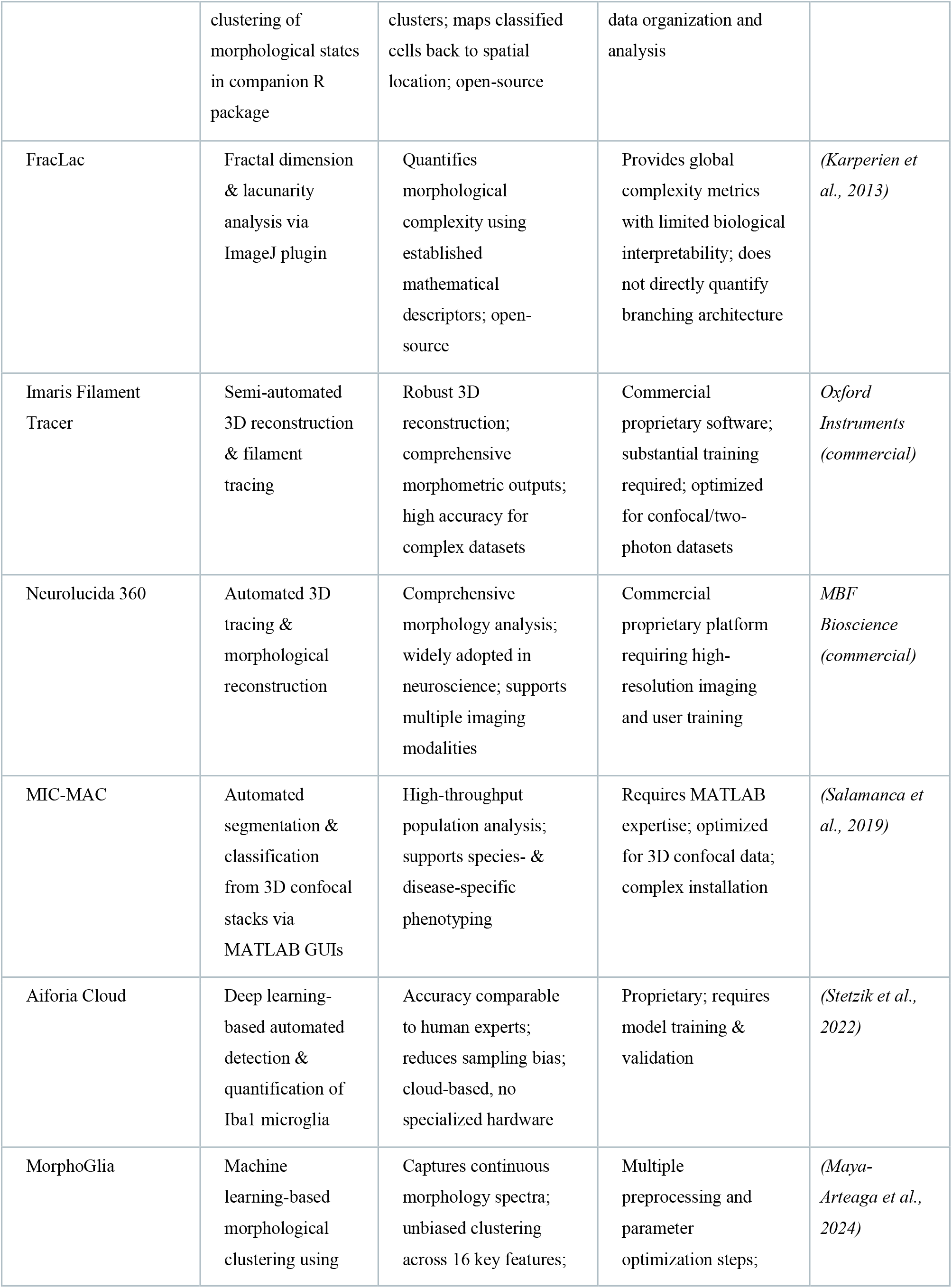
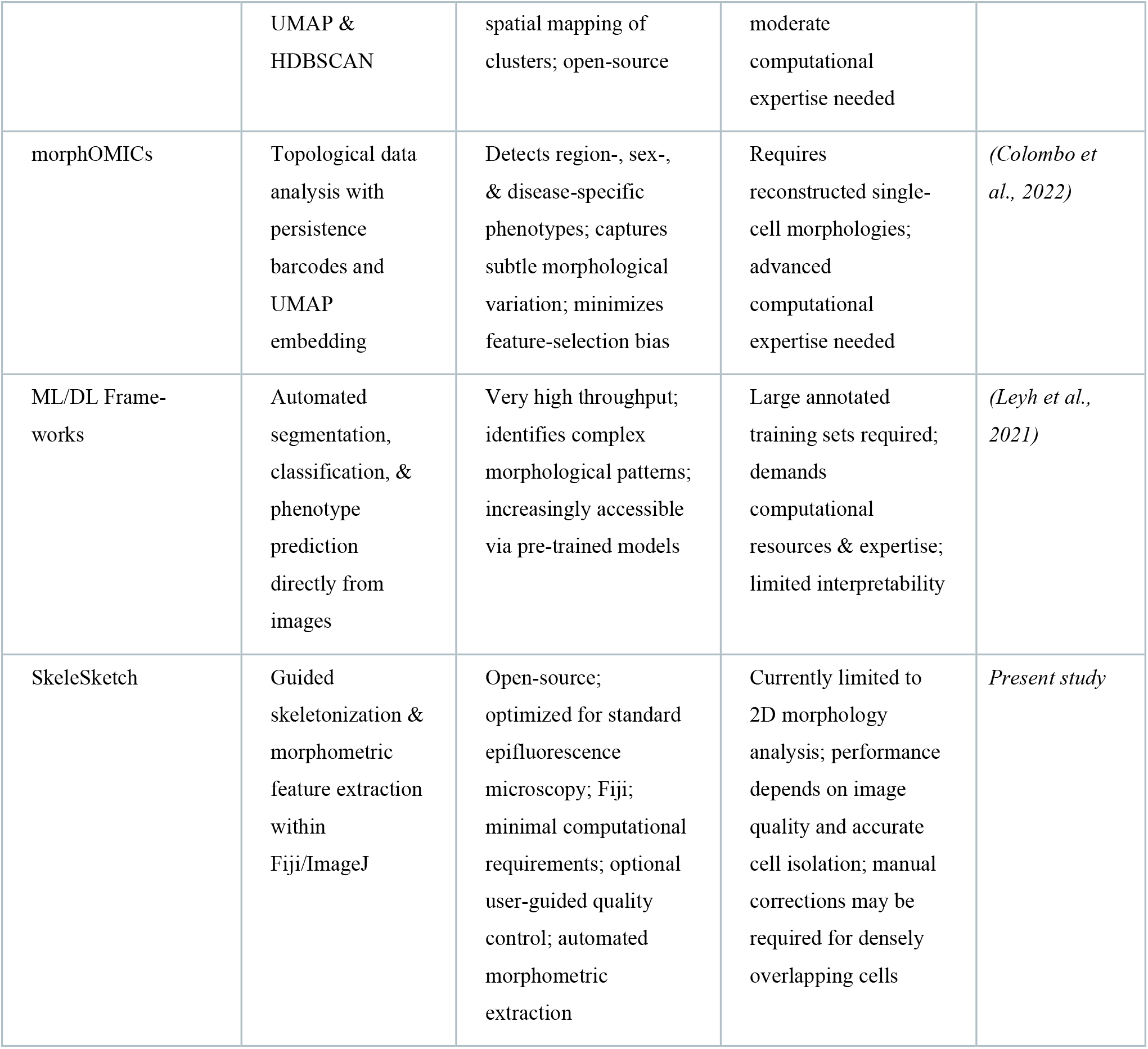
Comparison of tools and methods for quantitative microglial morphology analysis. For each tool, the primary analytical approach, principal strengths, principal limitations, and source reference are summarized. Tools span manual, semi-automated, and fully automated pipelines across open-source and proprietary platforms; SkeleSketch (present study) is included for comparison.

Here, SkeleSketch was developed as an open-source Fiji/ImageJ plugin for quantitative microglial morphology analysis from standard epifluorescence images. By integrating semi-automated skeletonization, user-guided quality control, and morphometric extraction into a single workflow, SkeleSketch provides an accessible alternative to labor-intensive manual tracing and computationally demanding image-analysis platforms. The plugin is freely available, easy to install, and requires only standard computing resources and minimal technical expertise. Using an established lipopolysaccharide (LPS)-induced neuroinflammation model, we demonstrate that our pipeline accurately reconstructs microglial morphology, captures the expected LPS-induced morphological changes, and generates concordant measurements across two commonly used labeling strategies (CX3CR1-GFP and Iba1) and the commercial platform, Imaris (Fig. 1).

**Figure 1.**
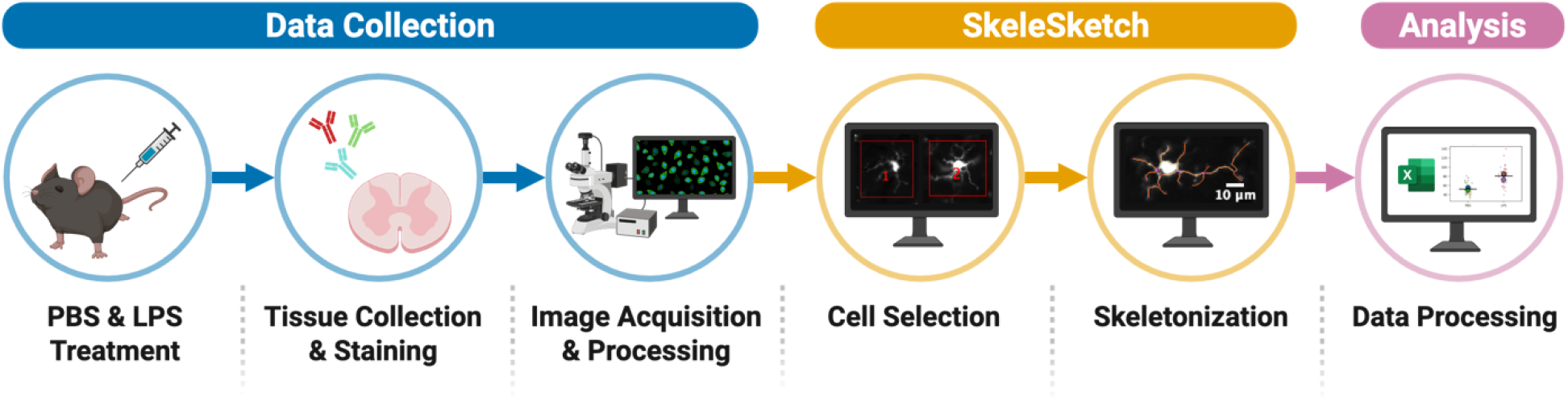
Study overview. Following PBS or LPS administration, spinal cord tissue was collected, immunostained, and imaged using standard epifluorescence microscopy. Individual microglia were selected and analyzed with SkeleSketch, which performed semi-automated skeletonization, morphometric feature extraction, and automated data export for downstream statistical analyses. Created in BioRender. Pascoe, L. (2026) https://BioRender.com/ycmqlyn

## Methods

### Animals

All mice were group-housed under standard conditions in the Van Andel Institute Vivarium and maintained on a 12h light/12h dark cycle, with lights on at ZT0 (7:00 A.M.). Experiments were conducted at approximately ZT2-ZT3. The CX3CR1-GFP knock-in mouse line was purchased from JAX (B6.129P2(Cg)-Cx3cr1tm1Litt/J, Cat. No. 005582), which expresses EGFP in monocytes, dendritic cells, NK cells, and brain microglia under control of the endogenous Cx3cr1 locus (Jung et al., 2000). Each treatment group consisted of three mice (two females and one male, 5–6 months of age). All procedures were approved by the Van Andel Institute Institutional Animal Care and Use Committee (IACUC) and conducted in accordance with institutional and NIH guidelines.

### LPS Treatment

Lipopolysaccharide (LPS; *E. coli* O55:B5, Sigma-Aldrich, Cat. No. 2637) was dissolved in PBS and administered intraperitoneally at 1 mg/kg every 24h for two consecutive days. PBS was used as the vehicle control and administered using identical parameters.

### Tissue Collection

Mice were killed 24h after the final injection via sodium pentobarbital overdose and transcardially perfused with 4% paraformaldehyde in PBS. Spinal cords were extracted and post-fixed in 4% paraformaldehyde for 24h, followed by cryoprotection in 30% sucrose for an additional 24h. Tissue was sectioned transversely at 20 µm using a Leica cryostat and collected sequentially.

### Immunofluorescence

Spinal cord sections (20 µm) were permeabilized in 1× PBS–Triton X-100 (0.3%) for 10 minutes and blocked with 1% BSA in PBS–Triton X-100 (0.3%) for one hour. Sections were then incubated overnight at 4°C with Rabbit anti-Iba1 (FUJIFILM Wako, Cat. No. 019-19741; 1:500). Following primary incubation, sections were washed three times in PBS–Tween 20 (0.1%; 5 minutes per wash) and incubated with the appropriate secondary antibody, Cy3 AffiniPure Goat Anti-Rabbit IgG (Jackson ImmunoResearch Laboratories, Cat. No. 111-165-144; 1:200). Sections were mounted using Fluoromount-G.

### Imaging

Sections of the ventral horn of lumbar spinal cord were imaged using an Olympus DP80 epifluorescence microscope at 20× magnification. Images were saved as individual 16-bit grayscale channels (FITC, TRITC, CY5, DAPI) via Olympus cellSens Dimension software, preserving full camera bit depth (14-bit) and embedded spatial calibration (0.51 μm/pixel at 20×). All images were acquired using identical microscope settings.

### Code Accessibility

The code/software described in the paper is freely available online at https://github.com/laurapascoe/Skelesketch and is archived on Zenodo (DOI: 10.5281/zenodo.22815758). A sample image is provided for testing. SkeleSketch was implemented in Fiji (ImageJ 1.54p; Schindelin et al., 2012) and run on both Windows 11 and macOS 26.2 using a standard desktop computer. Additional images used for the analyses in this paper are available from the authors upon request.

### Software Installation

ImageJ or Fiji can be installed from https://imagej.net/ij/download.html or https://fiji.sc/, respectively. Required dependencies (ImageJ, Fiji, Java-8, and ResultsToExcel) were added via Help > Update > Manage Update Sites. SkeleSketch was downloaded and placed in the Fiji plugins directory (Fiji.app > scripts > Plugins), ImageJ/Fiji was restarted to complete installation, and installation was verified by running the README via Plugins > CustomMacros. The README also documented the metrics captured, tunable parameters, and troubleshooting advice.

### Image Preparation

Images were loaded via File > Import > Bio-Formats. SkeleSketch requires grayscale images with a dark background relative to the foreground signal; images were converted to grayscale (Image > Lookup Table > Grays), adjusted to 16-bit depth (Image > Type > 16-bit), and background was inverted as needed (Image > Lookup Table > Invert LUT).

### Cell Selection

The prepared image was opened in ImageJ/Fiji and the F1 key was pressed to initiate cell selection (Extended Data Fig. 2-1). An output folder was specified, and individual microglia were selected with the polygon tool; the spacebar captured each selection, which was automatically labeled on the base image. After all cells were selected, Shift finalized the selection and opened the output folder. Only cells meeting predefined quality criteria for morphological analysis (i.e., complete soma visibility, sufficient separation from neighboring cells, adequate contrast between cell and background, full inclusion of the cell within the field of view) were selected.

### Skeletonization

The first cell image was opened from the output folder and F2 was pressed to begin skeletonization. Skeletons were manually corrected by setting the foreground color (Edit > Options > Colors) to black to remove excess branches or to white to add missing branches. The spacebar finalized edits to each skeleton, and Shift saved satisfactory results. The same mechanism applied to manual soma edits, which immediately followed the saved skeleton. Original images, skeletons, and composites were saved automatically to the output folder alongside an Excel file of cell morphology data. Subsequent images opened automatically, allowing continuous batch processing.

### Morphological Analysis

To assess SkeleSketch’s ability to detect expected treatment-associated morphological differences, a pilot validation dataset consisting of 3 PBS- and 3 LPS-treated animals was examined. LPS treatment produces a well-established phenotype characterized by soma enlargement, process retraction, and reduced branching complexity, providing a biologically relevant benchmark for evaluating the pipeline (Nakamura et al., 1999; Kloss et al., 2001; Melief et al., 2012). Twelve microglia were analyzed per animal by sampling three cells from each of four non-overlapping fields of view to capture within-animal morphological heterogeneity while minimizing sampling bias. Additionally, to further validate SkeleSketch, we evaluated its robustness across two commonly used microglial labeling strategies (GFP and Iba1) and benchmarked its morphometric outputs against those generated by Imaris.

SkeleSketch quantified a total of 18 morphometric features describing soma morphology, branching architecture, and process complexity. Because the importance of individual morphometrics varies depending on the biological question, users may select metrics appropriate for their experimental design. For our validation study, we focused on five representative measures that represent complementary aspects of microglial morphology and are widely reported in the literature: ramification index (total branch length / soma area), complexity index ((number of endpoints × total branch length) / soma area), soma area, total branch length, and maximum branch length. Together, these metrics characterize changes in cell size, process extension, and branching complexity that are expected to distinguish between PBS- and LPS-treated cells.

### Statistical Analysis

All analyses and figures were generated in Python (v3.11) using NumPy (v2.5.1), pandas (v3.0.4), SciPy (v1.18.0), and Matplotlib (v3.11.0). Because treatment was administered at the level of the animal, cell-level morphometrics were first aggregated to a single mean value per animal, and all between-group comparisons were performed on these animal-level summaries (n = 3 animals per group) to avoid pseudoreplication. Given the intentionally small, proof-of-principle cohort, group differences are reported descriptively rather than by null-hypothesis significance testing. For each metric, we report the group mean ± standard deviation, the difference in animal-level means (LPS − PBS, Δ), and its 95% confidence interval, calculated from the animal-level values using Welch’s t-distribution. Distributions are displayed as SuperPlots showing individual cell values, per-animal means, and the group mean ± standard error of the mean computed across animal means (Lord et al., 2020).

For the marker- and platform-concordance analyses, the same cells were measured under both conditions (CX3CR1-GFP vs Iba1; SkeleSketch vs Imaris Filament Tracer) and paired at the single-cell level. Agreement between paired measurements was quantified per metric using Pearson’s correlation coefficient (r) and Lin’s concordance correlation coefficient (CCC), the latter of which additionally penalizes systematic bias between methods. 95% confidence intervals for r and CCC were estimated by bias-corrected and accelerated bootstrap resampling (10,000 resamples). Composite indices (ramification and complexity) were recomputed from the corresponding Imaris primitives using the SkeleSketch definitions to ensure like-for-like comparison. Bracketed values represent the 95% confidence intervals throughout the Results.

## Results

### SkeleSketch reconstructs microglial morphology from standard epifluorescence images

Across both PBS- and LPS-treated samples, the pipeline consistently and reliably isolated individual microglia, skeletonized their processes, and segmented the soma. Optional user-guided correction of skeleton branches and soma masks was performed where automated segmentation was affected by uneven staining or intersecting processes (Fig. 2). PBS-treated microglia displayed the expected highly ramified morphology with long, fine processes and small soma, whereas LPS-treated microglia exhibited retracted, less complex branches and enlarged soma (Fig. 2) (Nakamura et al., 1999; Kloss et al., 2001; Melief et al., 2012).

**Figure 2.**
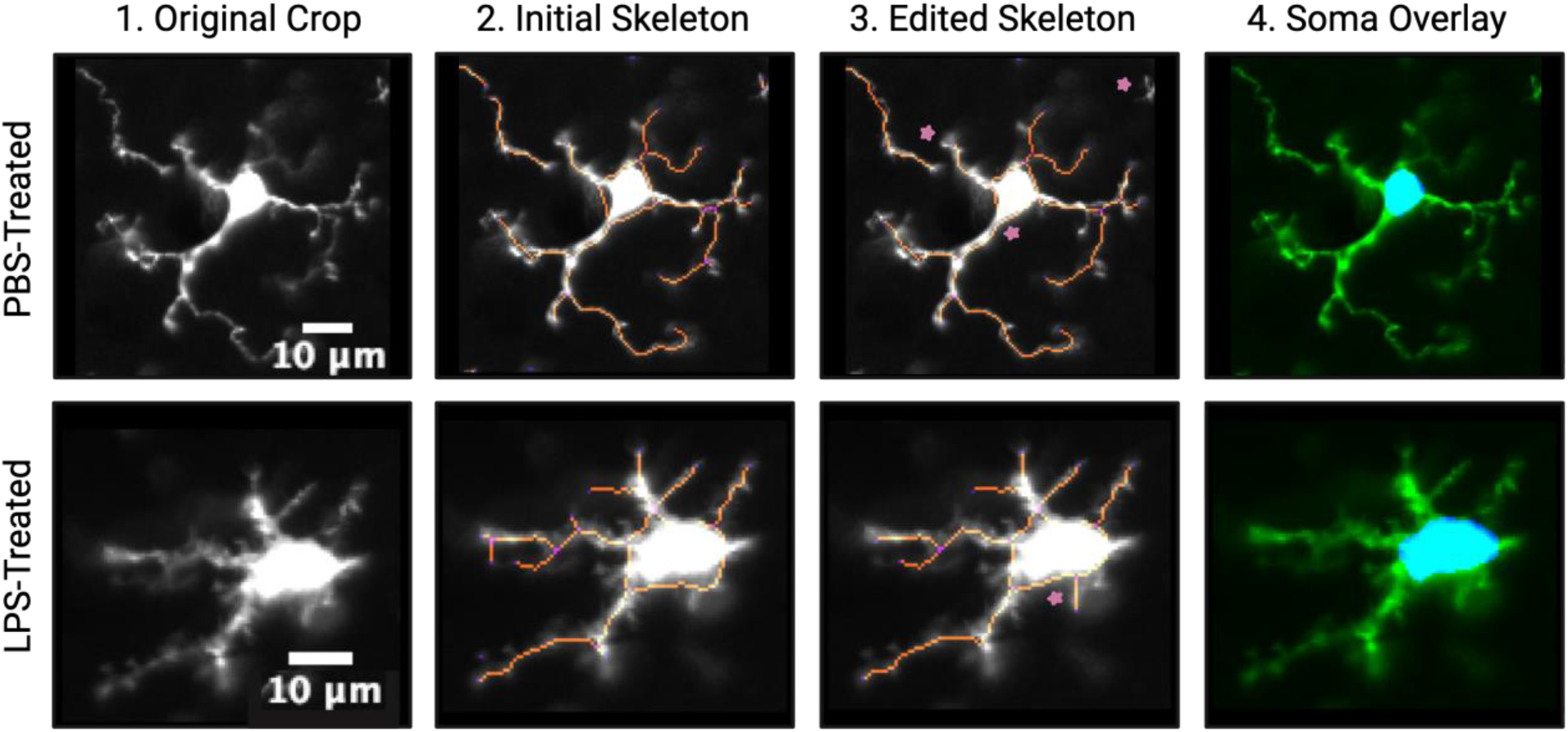
Representative reconstruction of spinal cord microglia by SkeleSketch. Representative spinal cord microglia from PBS-treated (top) and LPS-treated (bottom) mice are shown as the original cropped image (1), the initial, automatically generated skeleton (2), the manually corrected skeleton with edited regions indicated by stars (3), and the final soma segmentation overlay (4). Scale bar, 10 µm.

### SkeleSketch captures expected LPS-induced morphological remodeling

To evaluate whether the extracted morphometric measurements captured biologically meaningful variation, we compared morphological differences between PBS- and LPS-treated mice. For each metric, SuperPlots display the full distribution of cell-level values (small points), the corresponding animal means (large points), and the group mean (Fig. 3).

**Figure 3.**
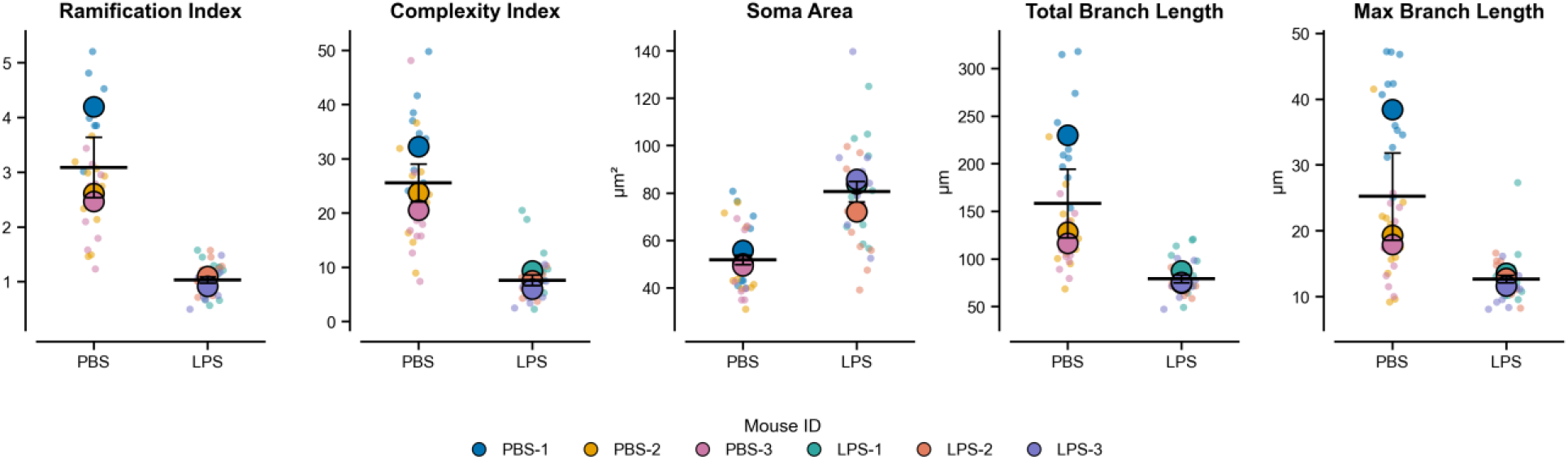
Comparison of five morphometrics between PBS- and LPS-treated groups. SuperPlots of commonly used morphometric measurements (left to right): ramification index (total branch length / soma area), complexity index ((number of endpoints × total branch length) / soma area), soma area, total branch length, and maximum branch length. Small dots represent individual cells and large dots represent per-animal means, colored by animal (mouse ID); horizontal bars and whiskers represent the group mean ± SEM computed across animal means (n = 3 animals per group).

SkeleSketch recovered the morphological changes classically associated with LPS treatment across all representative morphometric features (Fig. 3 and Extended Fig. 3-1). Ramification index (PBS 3.09 ± 0.96 vs LPS 1.03 ± 0.10; Δ = −2.06 [−4.40, 0.29])^a^, complexity index (PBS 25.5 ± 6.0 vs LPS 7.6 ± 1.7; Δ = −17.9 [−31.6, −4.2])^b^, total branch length (PBS 158.2 ± 62.5 µm vs LPS 79.0 ± 7.2 µm; Δ = −79.2 µm [−231.7, 73.4])^c^, and maximum branch length (PBS 25.2 ± 11.5 µm vs LPS 12.7 ± 1.0 µm; Δ = −12.6 µm [−40.7, 15.6])^d^ were each lower following LPS treatment. In contrast, soma area was greater in LPS-treated animals (PBS 51.8 ± 3.5 µm^2^ vs LPS 80.6 ± 7.4 µm^2^; Δ = +28.8 µm^2^ [13.3, 44.3])^e^. The largest and most consistent changes were in the complexity index and soma area; the wider intervals for the branch-length and ramification metrics reflect greater between-animal variability within the PBS group at this pilot cohort size, rather than inconsistent software performance. Similar trends were observed across the complete panel of extracted morphometrics (Extended Data Fig. 3-1), with the greatest separation for complexity- and branching-related measurements and minimal change in soma-shape descriptors (circularity, solidity, aspect ratio). Together, these findings indicate that SkeleSketch detects specific rather than uniform differences between groups.

### Morphometric readouts are concordant between GFP and Iba1 labeling

To determine whether SkeleSketch performs consistently across different labeling strategies, we compared morphometrics extracted from the same ten microglia labeled by both endogenous CX3CR1-GFP and Iba1 immunostaining. This matched-cell comparison was performed as a pilot in one PBS and one LPS animal. Accordingly, we interpret it as evidence of per-cell measurement consistency across labeling chemistries rather than a population-level claim across animals. Per-cell measurements showed strong agreement between the two markers across all five focus metrics: complexity index (r = 0.97 [0.87, 0.99], CCC = 0.82 [0.69, 0.95]; GFP 14.1 vs Iba1 19.7)^f^, total branch length (r = 0.93 [0.77, 0.98], CCC = 0.90 [0.82, 0.98]; GFP 105.1 vs Iba1 107.4 µm)^g^, ramification index (r = 0.88 [0.62, 0.93], CCC = 0.65 [0.34, 0.90]; GFP 1.4 vs Iba1 1.8)^h^, soma area (r = 0.77 [0.32, 0.95], CCC = 0.76 [0.52, 0.91]; GFP 93.1 vs Iba1 85.9 µm^2^)^i^, and maximum branch length (r = 0.76 [−0.03, 0.96], CCC = 0.68 [0.39, 0.97]; GFP 14.6 vs Iba1 13.5 µm)^j^. Both labeling strategies preserved the expected treatment-dependent differences in ramification and branching complexity, and PBS-treated microglia exhibited higher values than LPS-treated microglia (Fig. 4). Small systematic offsets were observed for some features. For example, Iba1 tended to yield slightly higher values for fine-branching metrics, consistent with differences in how the two markers label fine processes, but the relative differences between conditions were maintained across markers. Together, these results demonstrate that SkeleSketch generates robust morphometric measurements across both genetically encoded and immunohistochemical labeling strategies while preserving treatment-dependent morphological differences, supporting its application across diverse experimental models.

**Figure 4.**
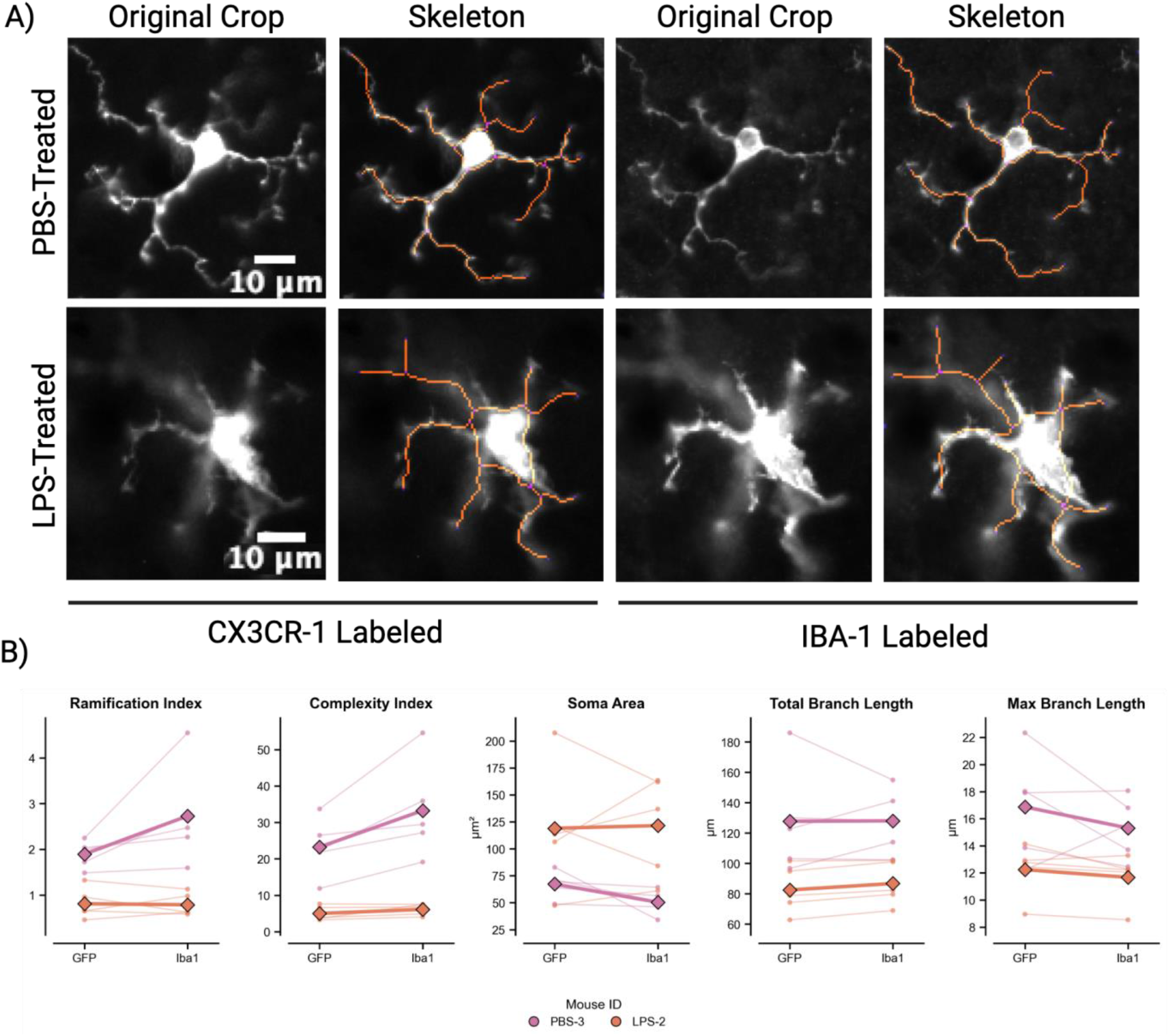
Validation of SkeleSketch across microglial labeling strategies. (A) Representative spinal cord microglia from PBS-treated (top) and LPS-treated (bottom) mice labeled with CX3CR1-GFP (left) or Iba1 immunostaining (right), with SkeleSketch-generated skeletons overlaid on the original fluorescence images. (B) Paired slope plots comparing morphometric measurements obtained from matched GFP- and Iba1-labeled cells (n = 10 matched cells). Each thin line represents a single matched cell, with connected points indicating paired measurements; bold lines represent group means, colored by animal (mouse ID). Scale bar, 10 µm.

### SkeleSketch measurements are concordant with Imaris Filament Tracer

To assess whether the morphometrics extracted by SkeleSketch are robust to the choice of analysis platform, we compared the same microglia analyzed independently by SkeleSketch and by Imaris Filament Tracer. Eighteen cells (three cells from each of three PBS- and three LPS-treated animals; representative cells skeletonized by both software packages (Fig. 5A)) were traced in both programs, and per-cell values for the five focus metrics were paired (Fig. 5B).

**Figure 5.**
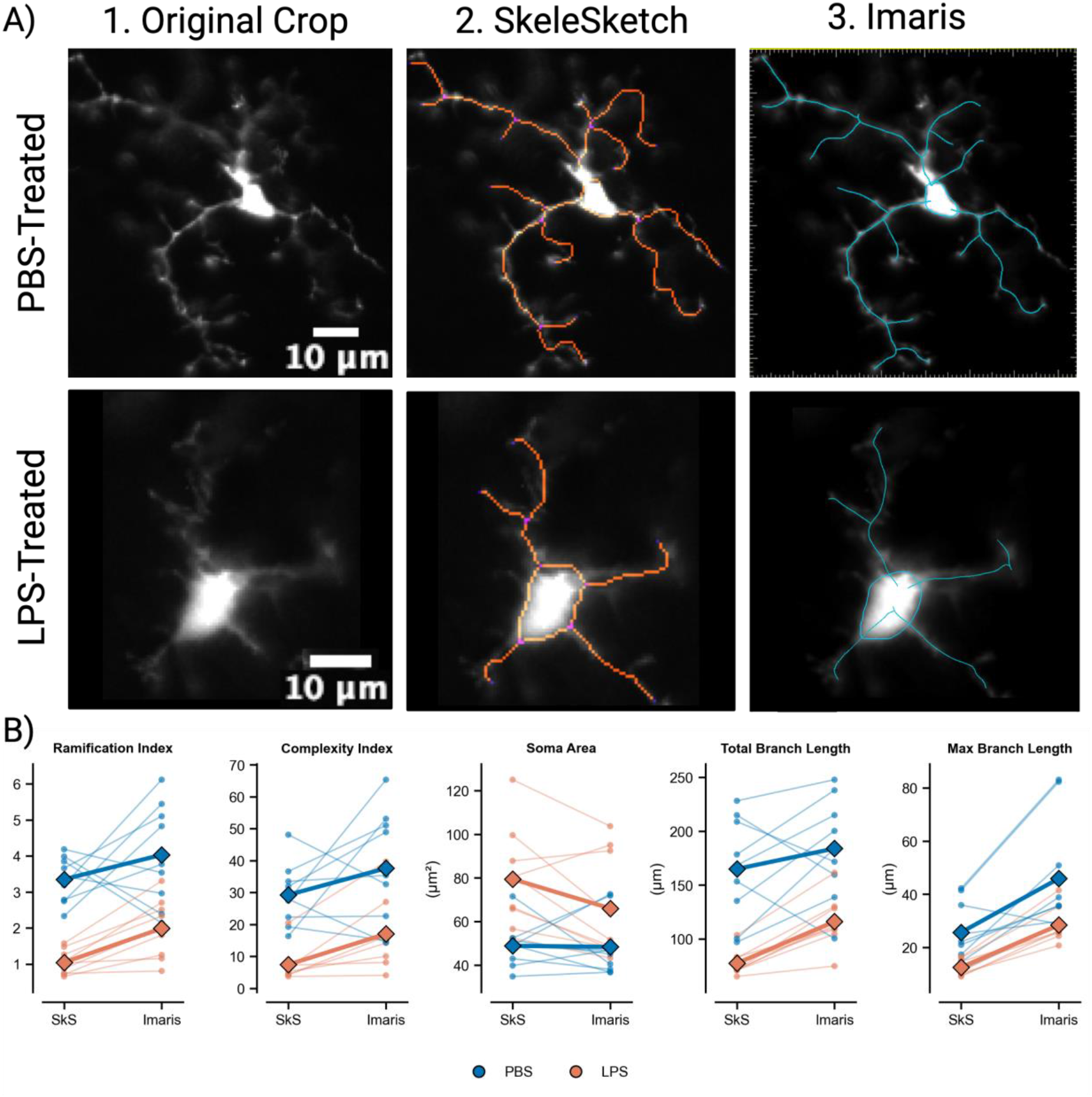
SkeleSketch morphometrics are concordant with Imaris Filament Tracer. (A) Representative microglia reconstructed independently by SkeleSketch and by Imaris Filament Tracer. (B) Paired slope plots comparing per-cell measurements of the five focus metrics obtained from the same microglia analyzed by both pipelines (n = 18 cells; three cells from each of three PBS- and three LPS-treated animals). Each thin line represents one cell, connecting its SkeleSketch (SkS) and Imaris value; bold lines represent group means; cells are colored by treatment group (PBS, blue; LPS, orange). Scale bar, 10 µm.

Across all five metrics, per-cell measurements were positively correlated between the two pipelines, with the closest agreement for the size- and length-based measures. Soma area showed the strongest concordance (r = 0.71 [0.37, 0.91], CCC = 0.67 [0.37, 0.88]; Imaris 57.2 vs SkeleSketch 64.2 µm^2^)^k^, and total branch length was correlated with a consistent offset (r = 0.73 [0.45, 0.92], CCC = 0.63 [0.43, 0.81]; Imaris 150.0 vs SkeleSketch 121.2 µm)^l^. Maximum branch length was strongly correlated but with a larger absolute offset (r = 0.78 [0.16, 0.97], CCC = 0.38 [0.09, 0.50]; Imaris 37.2 vs SkeleSketch 19.1 µm)^m^, as were the composite ramification (r = 0.57 [0.20, 0.79], CCC = 0.47 [0.25, 0.66])^n^ and complexity (r = 0.61 [0.29, 0.81], CCC = 0.50 [0.26, 0.71])° indices. The systematic offsets are attributable to well-defined methodological differences between the two platforms rather than measurement error. Imaris fits a smooth spline centerline whereas SkeleSketch measures along a pixelated skeleton, yielding consistently longer Imaris lengths. Imaris represents each process as a single soma-to-tip segment whereas SkeleSketch subdivides at every skeleton junction, inflating Imaris’ maximum branch length. Critically, both pipelines recovered the same direction of LPS-induced change for all five metrics, with PBS cells reading higher than LPS cells for ramification, complexity, total branch length, and maximum branch length, and lower for soma area. These results indicate that although absolute values differ between platforms in predictable ways, the biological conclusions drawn from SkeleSketch are preserved across analysis pipelines. As this comparison reflects agreement between two analytical platforms rather than accuracy against a ground-truth reference and was performed on a pilot set of 18 cells, the correlation values should be interpreted as preliminary. Full Imaris benchmarking parameters are provided in Extended Methods 1.

#### Statistical Table

Statistical analyses corresponding to values reported in the Results. Each superscript letter (a–o) in the text keys to a row, indicating the data structure and inferential method for that value. Rows a–e are animal-level comparisons between treatment groups (LPS − PBS); rows f–j and k–o are per-cell concordance analyses between labeling strategies (GFP vs Iba1) and between analysis platforms (SkeleSketch vs Imaris), respectively. r, Pearson’s correlation coefficient; CCC, Lin’s concordance correlation coefficient; CI, confidence interval. Bracketed values are 95% CIs (Welch’s t-distribution for animal-level differences; bias-corrected and accelerated bootstrap, 10,000 resamples, for r and CCC).

|  | Data structure | Type of test | 95% CI |
| --- | --- | --- | --- |
| a | Ramification index; animal-level means (n = 3/group) | Welch's t-based CI of difference in animal means (LPS – PBS) | [−4.40, 0.29] |
| b | Complexity index; animal-level means (n = 3/group) | Welch's t-based CI of difference in animal means (LPS – PBS) | [−31.6, −4.2] |
| c | Total branch length; animal-level means (n = 3/group) | Welch's t-based CI of difference in animal means (LPS – PBS) | [−231.7, 73.4] |
| d | Maximum branch length; animal-level means (n = 3/group) | Welch's t-based CI of difference in animal means (LPS – PBS) | [−40.7, 15.6] |
| e | Soma area; animal-level means (n = 3/group) | Welch's t-based CI of difference in animal means (LPS – PBS) | [13.3, 44.3] |
| f | Complexity index; paired per-cell, GFP vs Iba1 (n = 10 cells) | Pearson *r*; Lin's CCC | *r* [0.87, 0.99]; CCC [0.69, 0.95] |
| g | Total branch length; paired per-cell, GFP vs Iba1 (n = 10 cells) | Pearson *r*; Lin's CCC | *r* [0.77, 0.98]; CCC [0.82, 0.98] |
| h | Ramification index; paired per-cell, GFP vs Iba1 (n = 10 cells) | Pearson *r*; Lin's CCC | *r* [0.62, 0.93]; CCC [0.34, 0.90] |
| i | Soma area; paired per-cell, GFP vs Iba1 (n = 10 cells) | Pearson *r*; Lin's CCC | *r* [0.32, 0.95]; CCC [0.52, 0.91] |
| j | Maximum branch length; paired per-cell, GFP vs Iba1 (n = 10 cells) | Pearson *r*; Lin's CCC | *r* [−0.03, 0.96]; CCC [0.39, 0.97] |
| k | Soma area; paired per-cell, SkeleSketch vs Imaris (n = 18 cells) | Pearson *r*; Lin's CCC | *r* [0.37, 0.91]; CCC [0.37, 0.88] |
| l | Total branch length; paired per-cell, SkeleSketch vs Imaris (n = 18 cells) | Pearson *r*; Lin's CCC | *r* [0.45, 0.92]; CCC [0.43, 0.81] |
| m | Maximum branch length; paired per-cell, SkeleSketch vs Imaris (n = 18 cells) | Pearson *r*; Lin's CCC | *r* [0.16, 0.97]; CCC [0.09, 0.50] |
| n | Ramification index; paired per-cell, SkeleSketch vs Imaris (n = 18 cells) | Pearson *r*; Lin's CCC | *r* [0.20, 0.79]; CCC [0.25, 0.66] |
| o | Complexity index; paired per-cell, SkeleSketch vs Imaris (n = 18 cells) | Pearson *r*; Lin's CCC | *r* [0.29, 0.81]; CCC [0.26, 0.71] |

## Discussion

This study presents the initial validation of SkeleSketch, an open-source Fiji/ImageJ pipeline for quantitative analysis of microglial morphology from standard epifluorescence images. Using a pilot dataset of PBS- and LPS-treated mice, the pipeline captured the expected pattern of LPS-induced morphological remodeling (Nakamura et al., 1999; Kloss et al., 2001; Melief et al., 2012). Because LPS-induced morphological changes represent one of the most extensively characterized models of neuroinflammation, these findings provide proof-of-principle that SkeleSketch extracts biologically meaningful morphometric features.

Beyond biological sensitivity, SkeleSketch produced concordant measurements across both labeling strategies and analysis platforms tested. Morphometrics extracted from CX3CR1-GFP and Iba1-labeled cells showed strong per-cell agreement, and SkeleSketch’s outputs aligned with Imaris Filament Tracer in the direction of LPS-induced change across all five metrics. The observed offsets reflect known methodological differences in labeling chemistry and branch-length or soma-boundary definitions rather than measurement error, indicating that SkeleSketch performs reliably across common experimental variations in labeling and analysis software.

Morphometric analysis has become increasingly important as the field has moved away from labeling discrete microglial states. Capturing heterogeneity requires quantitative measurements that extend beyond categorical descriptors such as “ramified” or “amoeboid”. SkeleSketch was designed with this shift in mind, extracting continuous morphometric features at the single-cell level rather than assigning cells to predefined classes. This approach preserves biologically meaningful cell-to-cell variability, facilitates objective comparisons across experimental conditions, and enables detection of subtle structural changes that may be overlooked using categorical classification alone.

Existing methods for microglial morphology analysis provide powerful analytical capabilities but often require proprietary software, high-resolution imaging, complex computational workflows, or specialized user training (Table 1). These requirements can limit adoption across laboratories with varying imaging resources and computational expertise. SkeleSketch was developed to meet the need for an open-source workflow that integrates within an accessible platform (e.g., Fiji) and operates on standard epifluorescence images. The software requires only standard computing resources and minimal computational expertise, thus lowering the barrier to incorporating morphological measurements into routine workflows.

Another strength of the pipeline is the incorporation of optional user-guided refinement during skeletonization. Fully automated segmentation can struggle with uneven staining, intersecting processes, or imaging artifacts, particularly in lower-resolution datasets. Rather than eliminating user involvement entirely, SkeleSketch allows researchers to selectively and easily correct segmentation errors while preserving a largely automated workflow. This hybrid strategy provides flexibility across datasets that vary in staining intensity or image quality while substantially reducing the time and labor required in complete manual reconstruction. Collectively, these practical features broaden access to quantitative morphometric analysis and support its adoption across basic, translational, and educational neuroscience laboratories.

Several limitations should be considered in this study. First, the validation dataset was intentionally small and designed as a proof-of-principle rather than a comprehensive biological analysis. Larger validation cohorts spanning multiple brain regions, disease models, imaging systems, and independent laboratories will be important to further establish the robustness and reproducibility of the extracted morphometric features. Second, the current implementation is optimized for single-channel, two-dimensional epifluorescence images of individual cells and therefore requires adequate separation between neighboring microglia for accurate segmentation. Consequently, careful image acquisition and quality control remain important for reliable morphometric analysis. Third, because SkeleSketch allows for optional manual correction, its morphometric outputs may retain some operator bias. Formal assessment of inter-rater reliability, ideally under blinded conditions, would help establish the reproducibility of the correction step. Continued optimization of automated quality control, segmentation algorithms, and support for diverse imaging modalities will further improve robustness across diverse datasets. Because the pipeline is built entirely using open-source tools, it can be continuously refined and adapted as new methods become available. Future improvements could focus on automated cell detection, support for three-dimensional datasets, or untangling densely overlapping microglia. While this study focused on microglia, the underlying framework should be adaptable to other highly branched cellular populations. Neurons, astrocytes, oligodendrocyte precursor cells, and induced pluripotent stem cell-derived models all exhibit structural features that could be quantified using similar skeletonization-based approaches with appropriate optimization.

Overall, SkeleSketch provides an accessible and reproducible framework for quantitative microglial morphology analysis using widely available microscopy platforms and open-source software. By combining automated feature extraction with optional user-guided refinement, the pipeline enables objective characterization of microglial structure without requiring specialized imaging systems or advanced computational expertise. As morphometric analysis becomes increasingly important for understanding microglial biology, SkeleSketch provides a practical framework that can be readily adopted, independently validated, and further developed by the neuroscience community.

## Acknowledgements

During the preparation of this work, the authors used Claude (Anthropic) to assist with troubleshooting and debugging code for SkeleSketch, and ChatGPT (OpenAI) for grammar checking and language refinement of the paper. After using these tools, the authors reviewed and edited the content as needed and take full responsibility for the content of the publication.

## Author Contributions

LP, VM, SL, and QZ designed research, LP, VM, and SL performed research, LP and SL contributed unpublished reagents/ analytic tools, LP and VM analyzed data, LP, VM, and QZ wrote the paper.

