## Extended Data for "SkeleSketch: An Open-Source Tool for Quantifying Microglial Morphology in Fiji/ImageJ"

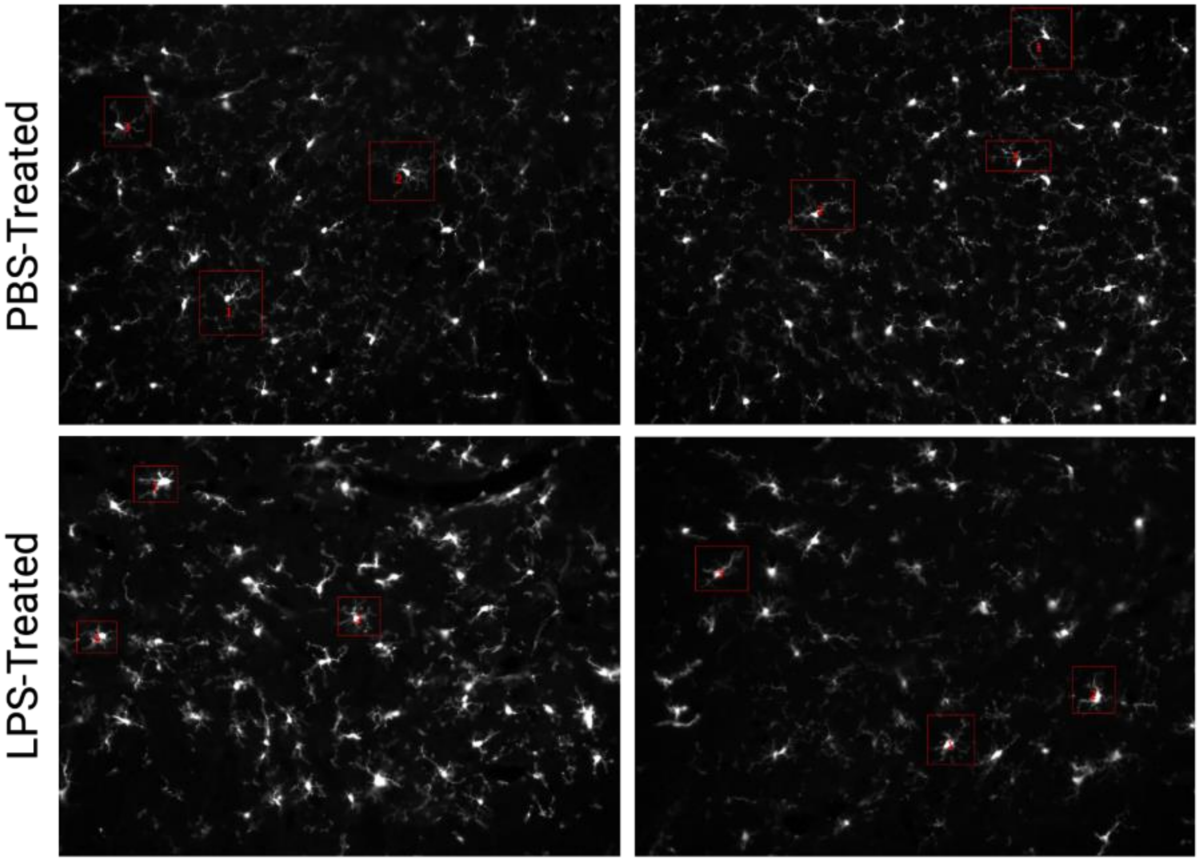


**Figure 2-1. Representative full-field images with labeled microglia selected for downstream analysis.** Full-field microglial images from PBS- and LPS-treated groups showing the cells selected for analysis (red boxes). This figure illustrates the image quality and cell-selection criteria required for SkeleSketch, including clearly defined soma, sufficient contrast between cell and background, and minimal overlap or intersection of processes with neighboring microglia to enable accurate segmentation and skeletonization. Extended Data for Figure 2.

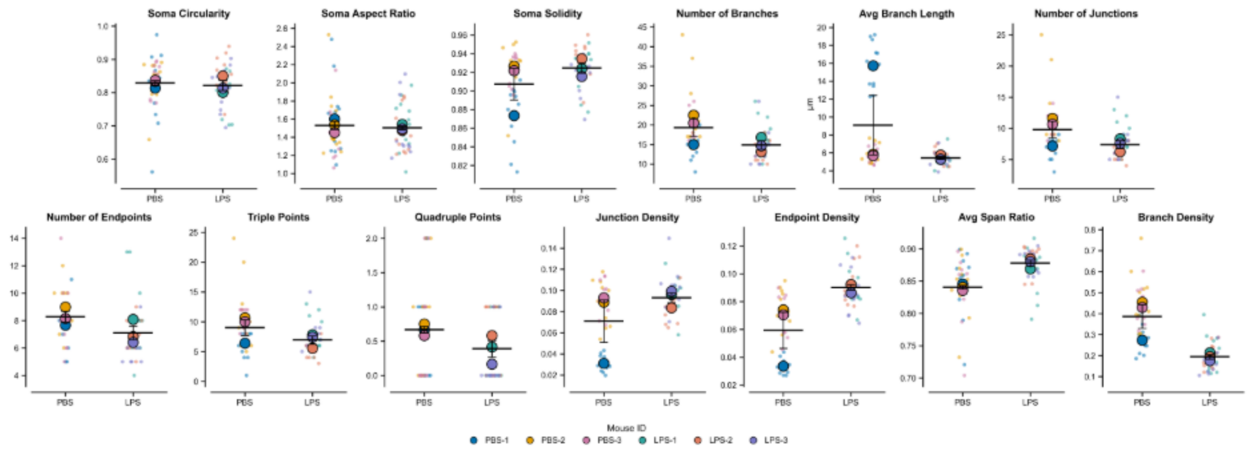


**Figure 3-1. Comparison of additional morphometric measurements between PBS- and LPS-treated groups.** SuperPlots of additional morphometric measurements generated by SkeleSketch, displayed as in Figure 3 (small dots, individual cells; large dots, per-animal means colored by mouse ID; bars, group mean ± SEM across animal means; n = 3 animals per group). Extended Data for Figure 3.

**Extended Methods 1. Detailed Imaris Filament Tracer benchmarking protocol.** Step-by-step Imaris workflow (v10.1.1, Oxford Instruments) used to independently reconstruct the 18 microglia (three cells from each of three PBS- and three LPS-treated animals) compared with SkeleSketch in Figure 5. The protocol covers image import and spatial calibration (0.5119 µm/pixel; two-dimensional analysis), machine-learning soma segmentation using the Surfaces object, AutoPath filament tracing with spine detection disabled, assignment of the segmented soma to the filament, and export of per-cell statistics with the mapping of Imaris outputs to the corresponding SkeleSketch metrics.

To assess concordance between SkeleSketch and an established commercial platform, the same cropped single-cell images analyzed by SkeleSketch were re-analyzed independently in Imaris (10.1.1, Oxford Instruments). Eighteen cells (three cells from each of three PBS- and three LPS-treated animals) were processed.

*Image Import*

Cropped single-cell TIFFs were collected into a single folder and opened in Imaris, converting each to the Imaris (.ims) format on import. Voxel size was confirmed under Edit → Image Properties to match the acquisition calibration (0.5119 µm/pixel), so that all measurements were reported in calibrated units (µm, $µm^2$). Images were analyzed in two dimensions.

*Soma Segmentation*

The soma was defined using the Surfaces object (Add New Surfaces). Default parameters were retained at the initial step, and the machine-learning segmentation mode was used; regions to retain were painted green and regions to exclude were painted pink. The machine-learning approach was selected because intensity-threshold methods failed to isolate the soma reliably in cells containing multiple bright regions. Segmentation was completed to generate a Surface corresponding to the soma.

*Filament Tracing*

Skeletons were generated using the Filaments object (Add New Filaments) with the AutoPath algorithm and spine detection disabled. The automated soma-detection steps of the wizard were advanced through, as the detected soma was subsequently replaced with the Surface generated above. At the seed-point step, Shift + left-click was used to place seed points at the tips of individual processes; AutoPath then traced each process from the soma to the corresponding seed point. Tracing was completed and filaments were displayed in line style.

*Soma Assignment*

On the Filament Edit tab, the wizard-generated soma was deleted and the previously created soma Surface was imported as the filament soma (Edit → Import Soma), so that all branch lengths were measured from the soma border and soma area was derived from the Surface.

*Data Export and Metric Mapping*

Per-cell statistics were exported from the Statistics tab (Detailed → export to .csv). The following Imaris outputs were mapped to the SkeleSketch metrics: Filament Length (sum) → total branch length; the maximum segment length → maximum branch length; Filament No. Segment Branch Points → junctions; Filament No. Segment Terminal Points → endpoints; and Soma Area → soma area. The composite ramification index (total branch length / soma area) and complexity index ((endpoints × total branch length) / soma area) were recomputed from these Imaris primitives using the same definitions applied by SkeleSketch, ensuring like-for-like comparison. Per-cell values from the two pipelines were paired and agreement quantified using Pearson's correlation coefficient and Lin's concordance correlation coefficient.
